# Single-nuclear RNA sequencing of endomyocardial biopsies identifies persistence of donor-recipient chimerism with distinct signatures in severe cardiac allograft vasculopathy

**DOI:** 10.1101/2022.08.24.504346

**Authors:** Kaushik Amancherla, Juan Qin, Michelle L Hulke, Ryan D Pfeiffer, Vineet Agrawal, Quanhu Sheng, Yaomin Xu, Kelly H Schlendorf, JoAnn Lindenfeld, Ravi V Shah, Jane E Freedman, Nathan R Tucker, Javid Moslehi

## Abstract

Cardiac allograft vasculopathy (CAV) is the leading cause of late allograft failure and mortality after heart transplantation. As current standards of diagnosis and treatment of CAV have significant limitations, understanding cell-specific responses may prove critical for developing improved detection strategies and novel therapeutics. This study is the first to successfully utilize human endomyocardial biopsy (EMB) samples to isolate large numbers of intact nuclei for single-nuclear transcriptomics. These data also lay the groundwork for ongoing experiments to study serial, routinely-collected EMB specimens after heart transplantation to identify novel biomarkers and pathways through which early CAV pathogenesis can be interrupted, thereby prolonging allograft survival.

## INTRODUCTION

Cardiac allograft vasculopathy (CAV) is the leading cause of late allograft failure and mortality after heart transplantation^1^. Histologically, CAV is chronic vascular rejection characterized by diffuse intimal thickening of macro- and microvasculature. While *in vitro* cellular models and *in vivo* histologic observations suggest coordinated responses of endothelial, fibroblast, and smooth muscle cells in CAV pathology, cell-specific transcriptional signatures among these in the transplanted human heart have not been studied. As current standards of diagnosis and treatment of CAV have significant limitations, understanding cell-specific responses may prove critical for developing improved detection strategies and novel therapeutics.

Here, we used single-nuclear RNA sequencing (snRNA-seq) to elucidate the transcriptomic landscape of CAV. Importantly, we establish the feasibility of performing snRNA-seq from human endomyocardial biopsy (EMB) specimens (3-10 mg in size) obtained at the time of right heart catheterization, enabling high-resolution molecular profiling of samples collected during routine clinical practice. We compared tissue obtained at the time of re-transplantation from 4 individuals with severe CAV to EMB specimens from 3 individuals post-transplant without CAV (**Figure 1A**). For all 7 individuals, samples were obtained from the right ventricle (RV). In 3 out of 4 patients with severe CAV, left ventricular (LV) samples were also obtained (for a total 10 samples).

**Figure 1.**
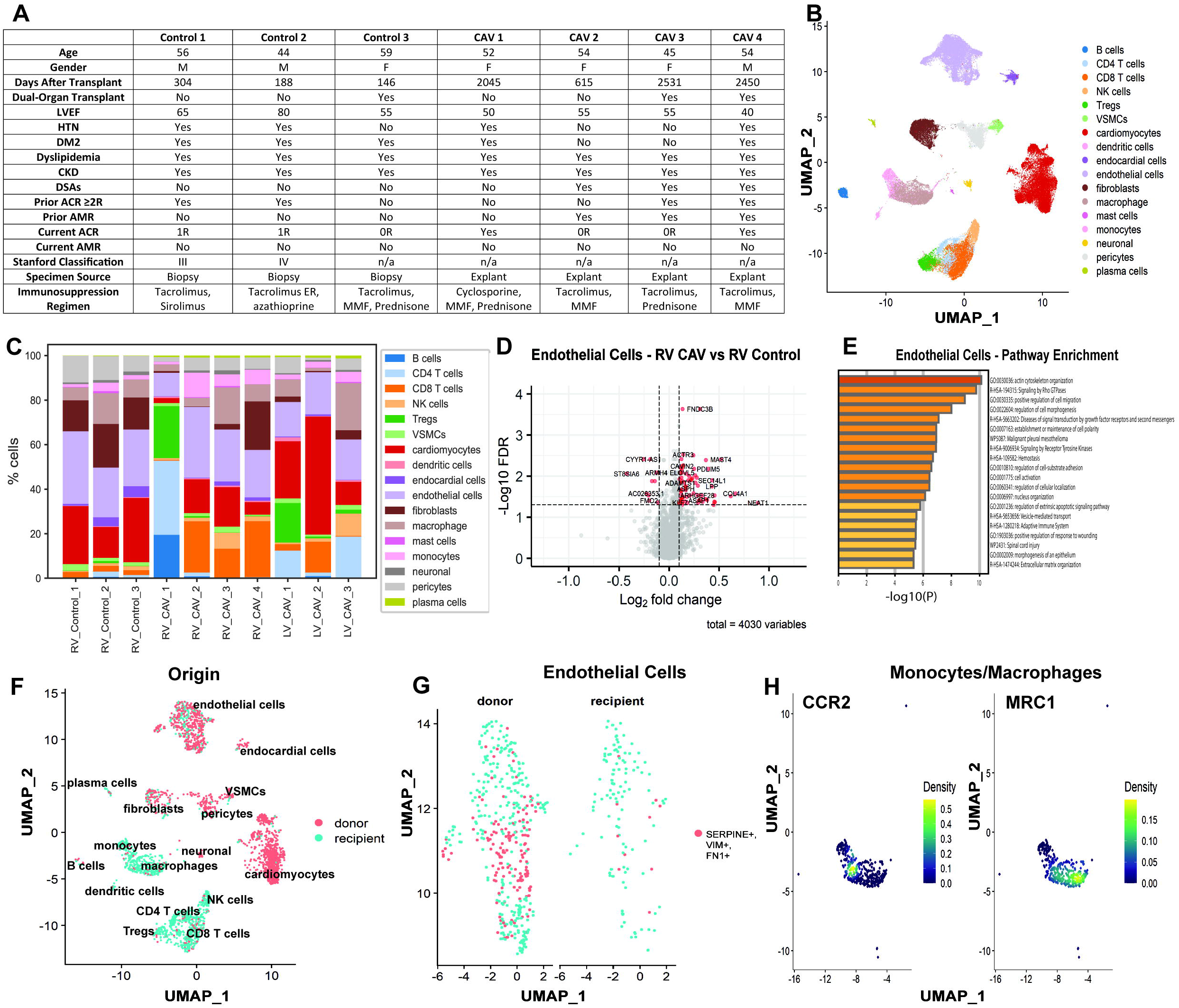
**A)** Clinical characteristics of all seven patients studied. **B)** Uniform Manifold Approximation and Projection (UMAP) of 62,465 nuclei identified 17 major cell types using canonical marker genes. **C)** Cell compositional analyses were performed using scCODA v0.1.6. Labels correspond to patients described in *Figure 1A*. No significant difference in cell composition was noted using an automated reference cluster. **D)** Differential gene expression was performed using MAST. Volcano plot representing right ventricular CAV vs. control samples for the endothelial cell cluster. **E)** Biological pathway enrichment analysis of differentially upregulated genes in RV CAV endothelial cells using Metascape. **F)** Genotype-free inference of donor-versus recipient-derived nuclei was performed using souporcell v2.0. UMAP of donor-vs. recipient-derived nuclei. The clusters correspond to the same clusters annotated in *Figure 1B*. **G)** Donor-derived endothelial cells are enriched for markers of endothelial-to-mesenchymal transition. **H)** The monocyte/macrophage cluster is largely recipient-derived. Presence of distinct *CCR2*^+^ monocytes and *CCR2*^-^*MCR1*^+^ macrophages is highlighted using Nebulosa. *LVEF = left ventricular ejection fraction; HTN = hypertension; DM2 = type 2 diabetes mellitus; CKD = chronic kidney disease; DSA = donor-specific antibodies; ACR = acute cellular rejection; AMR = antibody-mediated rejection; ER = extended release; MMF = mycophenolate mofetil; RV = samples from* right *ventricle; LV = samples from left ventricle; CAV = cardiac allograft vasculopathy; EndoMT = endothelial-to-mesenchymal transition*.

## METHODS

After nuclear isolation with modifications to account for low tissue mass, libraries were generated, sequenced, quality-controlled, and analyzed as previously described^2^. Raw FASTQ files are deposited at the NIH NCBI GEO data repository (GSE203548) and code used for these analyses are deposited at https://github.com/learning-MD/CAV. This study was approved by the Vanderbilt University Medical Center’s Institutional Review Board.

## RESULTS

We successfully isolated 62,465 nuclei and identified 17 major cell types with heterogenous distribution across the ten different samples (**Figures 1B, 1C**). When comparing RV samples, endothelial cells and fibroblasts in CAV exhibited increased expression of *SERPINE1*, which promotes neointimal hyperplasia and fibrosis^3^. Endothelial cells were enriched for pathways involved in angiogenesis, cell migration, and extracellular matrix (ECM) organization (**Figures 1D, 1E**). Fibroblasts in CAV exhibited increased expression of genes involved in ECM deposition and fibrosis (e.g., *MMP2, CCN1, THBS1*) while also highly expressing *IL6ST*, involved in IL-6 signaling. As expected, macrophages in CAV showed increased expression of genes associated with inflammation (e.g., *TLR2, IFNAR2*). While no significant differences in T cells were noted between conditions, subclusters included CD4 central memory T cells (*IL7R, TCF7*), CD4 T regulatory cells (*FOXP3, CTLA4, IL2RA*), and CD4 T cells exhibiting markers of exhaustion (*LAG3, CTLA4, PDCD1*), along with CD8 memory T cells (*CCL5*). No major differences in gene expression were noted between RV and LV CAV samples.

We repurposed a genotype-free demultiplexing tool to infer donor- and recipient-derived nuclei from each individual CAV sample. Using 5 of the 7 combined CAV samples (including both LV and RV tissue), 2,827 nuclei were confidently called as donor- or recipient-derived in the absence of genotyping (**Figure 1F**). Endothelial cells exhibited significant donor-recipient chimerism (21.8% recipient-derived). Donor-derived endothelial cells were enriched for markers of endothelial-to-mesenchymal transition (EndoMT; *SERPINE1, VIM, COL3A1*; **Figure 1G**). In contrast, immune cells were largely replaced by those originating from the recipient (91.1% of macrophages/monocytes, 92.6% of NK cells, 88% of T cells). Recipient-derived macrophages included both *CCR2*^*+*^ monocyte-derived macrophages and CCR2^-^ *MRC1*^+^ tissue resident macrophages, traditionally thought to be involved in cardiac repair^4^ (**Figure 1H**). Macrophages exhibited markers of activation, including *HLA-DRA* and *CD74*, and increased expression of *TGFB1*, a potential driver for the EndoMT observed in donor-derived endothelial cells.

## DISCUSSION

This study is the first to successfully utilize human EMB samples to isolate large numbers of intact nuclei for single-nuclear transcriptomics. As expected from an ischemic allograft, we see enrichment for genes and pathways involved in inflammation, fibrosis, and tissue healing. We highlight several unique findings enabled by this approach: 1) cell composition amongst EMB samples is highly heterogeneous, suggesting that bulk RNA-seq approaches may exhibit high levels of variability due to sampling bias; 2) there are unique transcriptomic signatures of donor-versus recipient-derived cells, particularly endothelial cells, highlighting putative novel avenues for investigation; and 3) the presence of recipient-derived CCR2^-^ macrophages warrants further study, as only a small percentage would be expected to be recipient-derived^4^. However, recent single-cell data have implicated partial replacement of MHC-II^hi^CCR2^-^ cardiac macrophages by monocytes, suggesting a still evolving understanding of macrophage subsets^5^.

Our study is limited by a small sample size and the use of samples derived from severe CAV. However, these data demonstrate feasibility of performing snRNA-seq using frozen EMBs, presenting a unique opportunity that may have broad ramifications on the fields of heart transplantation and cardio-oncology/immunology. These data also lay the groundwork for ongoing experiments to study serial, routinely-collected EMB specimens after heart transplantation to identify novel biomarkers and pathways through which early CAV pathogenesis can be interrupted, thereby prolonging allograft survival.

